# Impact of fenbendazole resistance in *Ascaridia dissimilis* on the economics of production in turkeys

**DOI:** 10.1101/2021.03.30.437784

**Authors:** James B. Collins, Brian Jordan, Anand N. Vidyashankar, Pablo Jimenez Castro, Ray M. Kaplan

## Abstract

Feed conversion efficiency is among the most important factors affecting profitable production of poultry. Infections with parasitic nematodes can decrease efficiency of production, making parasite control through the use of anthelmintics an important component of health management. In ruminants and horses, anthelmintic resistance is highly prevalent in many of the most important nematode species, which greatly impacts their control. Recently, we identified resistance to fenbendazole in an isolate of *Ascaridia dissimilis*, the most common intestinal helminth of turkeys. Using this drug-resistant isolate, we investigated the impact that failure to control infections has on weight gain and feed conversion in growing turkeys. Birds were infected on Day 0 with either a fenbendazole-susceptible or -resistant isolate, and then half were treated with fenbendazole (SafeGuard^®^ Aquasol) at 4- and 8-weeks post infection. Feed intake and bird weight were measured for each pen weekly throughout the study, and feed conversion rate was calculated. Necropsy was performed on birds from each treatment group to assess worm burdens at weeks 7 and 9 post infection. In the birds infected with the susceptible isolate, fenbendazole-treated groups had significantly better feed conversion as compared to untreated groups. In contrast, there were no significant differences in feed conversion between the fenbendazole-treated and untreated groups in the birds infected with the resistant isolate. At both weeks 7 and 9, worm burdens were significantly different between the treated and untreated birds infected with the drug-susceptible isolate, but not in the birds infected with the drug-resistant isolate. These significant effects on feed conversion were seen despite having a rather low worm establishment in the birds. Overall, these data indicate that *A. dissimilis* can produce significant reductions in feed conversion, and that failure of treatment due to the presence of fenbendazole-resistant worms can have a significant economic impact on turkey production. Furthermore, given the low worm burdens and an abbreviated grow out period of this study, the levels of production loss we measured may be an underestimate of the true impact that fenbendazole-resistant worms may have on a commercial operation.

## 1. INTRODUCTION

Both helminth and protozoan parasites can impact poultry performance parameters such as weight gain and/or feed conversion ratio (FCR) (Voeten, Braunius et al. 1988, Daş, Kaufmann et al. 2010, Sharma, Hunt et al. 2019). Feed conversion, a measure of feed consumption per unit of production accounts for approximately 70% of production costs, making it among the most important factors affecting profitable production (Willems, Miller et al. 2013). A lower feed conversion ratio (FCR) indicates that feed is being more efficiently utilized for growth. While coccidia (*Eimeria* spp.) are well documented as important parasitic pathogens of poultry, helminths generally receive much less attention. Several studies in chickens have shown that infections with *Ascaridia galli* have a negative impact on both feed efficiency and egg quality (Daş, Kaufmann et al. 2010, Stehr, Grashorn et al. 2019). However, less work has been done investigating this issue in turkeys infected with *Ascaridia dissimilis*.

*Ascaridia dissimilis* is the most prevalent and one of the most important parasites of turkeys, with up to 100% of a flock being infected (Yazwinski, Tucker et al. 2009). *Ascaridia* eggs are capable of surviving the environmental extremes that are present in poultry houses and may remain infective for periods exceeding six months, leading to a cycle of continuous reinfection and environmental contamination with new eggs (Cauthen 1931, Tarbiat, Jansson et al. 2015). Heavy infections may cause clinical disease such as diarrhea, intestinal blockage, and enteritis, but most often infections are subclinical, only causing reduced feed efficiency (Ikeme 1971, Norton, Hopkins et al. 1992, Yazwinski, Tucker et al. 2002). Given the potential health and production impacts of *Ascaridia*, as well as its near ubiquity, successful control will often be important for profitable production.

Currently, in the United States, fenbendazole is the only available treatment approved by the Food and Drug Administration for treatment of *Ascaridia* infections in poultry. Registration studies of fenbendazole (SafeGuard®) in feed, at 1mg/kg body weight for 6 days, demonstrated greater than 99% efficacy against *Ascaridia dissimilis* (United States Food and Drug Administration 2000). In addition, a formulation of fenbendazole that is administered in water, (SafeGuard^®^ Aquasol), demonstrated a mean efficacy of 97.7% against *Ascaridia galli*, a closely related parasite of chickens, that may also infect turkeys (United States Food and Drug Administration 2018). On commercial turkey farms, treatments with fenbendazole are often administered frequently, around every 4 weeks, which is an interval less that the prepatent period *A. dissimilis*. These treatments are typically administered in either feed or water to the entirety of the house. These means of drug delivery make accurate dosing challenging due to difficulty in optimal delivery of the drug and variability in consumption. Both issues may lead to sub-therapeutic levels of ingestion in some birds. In other livestock species, under-dosing is thought to be an important factor influencing the development of drug resistance in nematode parasites (Smith, Grenfell et al. 1999, Jackson and Coop 2000). A model investigating factors promoting the development of anthelmintic resistance showed that repeated under-dosing acted as a strong selector for resistance, since partially resistant heterozygotes were able to survive and reproduce (Smith, Grenfell et al. 1999). The survival of heterozygotes led to a much more rapid increase in the frequency of resistant homozygotes in the population as compared to full-dose treatments that killed the heterozygotes with high efficacy. Under-dosing, combined with often intensive use in production animals, may act as strong selectors for the development of anthelminthic resistance in nematode parasites.

In many species of important livestock parasites, resistance to benzimidazoles is highly prevalent (Kaplan 2004, Howell, Burke et al. 2008, Kaplan and Vidyashankar 2012). Though reduced efficacy of fenbendazole was reported previously in *Ascaridia dissimilis*, (Yazwinski, Tucker et al. 2013) resistance to fenbendazole in *A. dissimilis* was only recently confirmed for the first time in a controlled efficacy study (Collins, Jordan et al. 2019). Following treatment with fenbendazole, a field isolate of *A. dissimilis* (Sn) yielded an efficacy of 63.9%, whereas in three other field isolates fenbendazole treatment yielded an efficacy of greater than 99%. Having demonstrated fenbendazole resistance in a naturally occurring field isolate of *Ascaridia dissimilis,* we wanted to measure the effects that resistant parasites may be having on production parameters as a consequence of failed treatments.

## 2. MATERIALS AND METHODS

### 2.1 Turkeys and feeding

Four hundred and thirty-two, day old, Hybrid turkey poults were received from Prestage Farms and housed at the Poultry Science farm at the University of Georgia. Birds were allowed one week of acclimation before the study began. Water and feed were provided *ad libitum*. For the first 6 weeks, birds were fed a starter ration with 26% protein, then a grower ration with 23% protein was offered from weeks 6 to 9 (see Supplemental files 1 & 2 for the diet formulations).

### 2.2 Study Design

Birds were received on Day −7 and were assigned to 36 pens of 12 birds each based on weight, minimizing differences in total weight between pens. 16 pens were infected with the resistant isolate, 16 pens were infected with the susceptible isolate, and 4 pens served as environmental controls. Groups were separated by floor to ceiling mesh curtains to prevent movement of birds between pens. Feed was added into hanging feeders and the initial weight of feeders for each pen was recorded. Each subsequent week, total bird weight for each pen and the weights of feeders were recorded to determine the weight gain and feed consumed. The hanging feeders were then refilled and an initial feeder weight for the next week was recorded. At weeks 7 and 8 post infection (p.i.), groups were culled to 10 and 9 birds respectively, to maintain recommended stocking densities. The study was originally planned to continue for 16 weeks but was terminated at week 9 due to inability of the facilities to properly contain turkeys of this size. Birds were necropsied, and worm enumeration was performed on 8 and 16 birds for each treatment at weeks 7 and 9 p.i., respectively.

### 2.3 Parasite Isolates

Eggs from a resistant (Sn 3.1F2F) and a susceptible (Ow 3.0) isolate of *A. dissimilis* were obtained from passage of isolates whose drug susceptibility phenotypes were previously confirmed (Collins, Jordan et al. 2019). Briefly, feces containing *A. dissimilis* eggs were washed through a series of sieves, and then eggs were isolated by flotation using a solution with specific gravity of 1.15 and centrifuged at 433g for 7 mins. The supernatant was collected on a 32um mesh sieve and rinsed to remove flotation solution from eggs. Eggs were then stored in a tissue culture flask containing water and 0.5% formalin and stored at 25°C to allow development to the third stage larvae or infective stage.

### 2.4 Infection and Treatment

Starting on Day 0, 16 groups were infected with eggs of the resistant Sn 3.1F2F isolate (hereafter referred to as Sn) and 16 groups were infected with the susceptible Ow 3.0 isolate (hereafter referred to as Ow). Half of the groups infected with each isolate were then left untreated and half received treatment with fenbendazole at weeks 4 and 8 (p.i.). In addition, 4 groups of 12 birds each were included as uninfected environmental sentinels.

Each week, fully larvated infective *A. dissimilis* eggs were mixed into feed at a target inoculum dose of 25 eggs per bird. 3600 fully developed infective eggs in a volume of 1 ml were pipetted onto 360 grams of feed, and the feed was then mixed well to disperse the eggs. Twenty gr aliquots of the egg-contaminated feed containing approximately 300 eggs were then delivered to each group each week by sprinkling on top of the fresh feed, adjusting to 250 and 225 total eggs as birds were culled at weeks 7 and 8 p.i.

At weeks 4 and 8, treated groups were administered fenbendazole for five consecutive days at a dosage of 1.25 mg/kg, which is 25% higher than the recommended label dose of 1.0 mg/kg. This higher dose was provided to maximize the likelihood that all birds consumed the minimum full label dose. Treatment was administered using carboys delivering water to two side by side pens. Dosage was calculated based on the total bird weight for both pens, selected 1 day prior to the initiation of treatment. In order to maximize the likelihood that all birds would consume the full dosage, the fenbendazole was administered in 90% of the estimated volume of total daily water consumption. On all treatment days, the full volume of water containing the fenbendazole was consumed.

### 2.5 Statistical Analysis

Statistical analyses were performed on weight gain and FCR values to model and identify the effect of treatment, specifically, comparing turkeys infected with Ow and Sn, respectively. Data from both week 4 and week 5 was considered as baseline in separate analyses. To account for the growth across time, both linear and quadratic effects were introduced into the model. Likelihood based methods were used for statistical analyses.

Specifically, the fitted model for Weight gain data was:

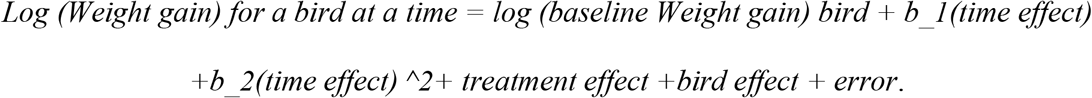

Conversely for FCR data was:

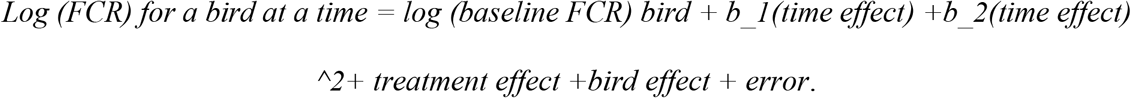

The error was assumed to be normally distributed with mean 0 and variance that changed with the treatment group. The errors between time points were modeled as an autoregressive model of order 1 that changed across treatment groups. The bird effect was treated as a random effect that was normally distributed with mean zero and independent of the error. All models were selected using the Bayesian Information Criterion after considering several polynomial models for time and different covariance structures. The normality of the error distributions was evaluated using Shapiro-Wilks test.

The number of immature and adult worms recovered on day seven and day nine was statistically analyzed, separately, using negative binomial regression with the logarithmic link function. This model was chosen based on the likelihood criterion. In the analyses for adult worms, data for the treated Ow group was not used in the analysis since all the observations were zero. The model included the treatment group as an effect. All statistical comparisons were evaluated at a 5% level of significance.

## 3. RESULTS

Analyses for weight gain and feed conversion ratio were performed separately using either week 4 or week 5 as baseline, with both analyses yielding consistent results. Week 5 was selected as the baseline for the results presented here, and results using week 4 as baseline are provided in Supplementary Tables 1 and 2.

### Weight Gain

Based on the fitted model, the distribution of the errors was found to be normal (p-value=0.0871), and baseline was not a significant factor (p-value=0.3843). The slope for week was estimated to be −0.1154 (Std. Error=0.0810), and the slope for the square of time was estimated to be 0.0238 (Std. Error=0.0160), both of which were not significantly different from zero (p-values=0.1406, 0.1574). Weight gains (Table 1) were not significantly different between experimental groups (p-value=0.1283).

**Table 1.** Weight Gain (kgs) for each treatment group by week. There were no significant differences in weight gain between the groups.

| Week | Treatment |  |  |  |
| --- | --- | --- | --- | --- |
|  | Ow-Treated | Ow-Untreated | Sn-Treated | Sn-Untreated |
| 1 | 0.14 | 0.15 | 0.16 | 0.16 |
| 2 | 0.20 | 0.21 | 0.17 | 0.21 |
| 3 | 0.33 | 0.36 | 0.34 | 0.35 |
| 4 | 0.43 | 0.45 | 0.45 | 0.44 |
| 5 | 0.44 | 0.54 | 0.48 | 0.49 |
| 6 | 0.70 | 0.63 | 0.71 | 0.65 |
| 7 | 0.77 | 0.79 | 0.80 | 0.83 |
| 8 | 0.56 | 0.70 | 0.61 | 0.64 |
| 9 | 0.72 | 0.77 | 0.89 | 0.76 |

### Feed Conversion Ratio

Based on the fitted model, the distribution of the errors was found to be normal (p-value=0.5040), and baseline was not a significant factor (p-value=0.6035). The slope for week was estimated to be 0.3571 (Std. Error= 0.0866) and slope for the square of time was estimated to be −0.0412 (Std. Error= 0.0171), both of which were significantly different from zero (p-values <0.0001, = 0.0179). Feed Conversion Ratio values are shown in Table 2 and Figure 1. Least square mean values for Feed Conversion Ratio (Table 3) differed overall between the groups (p-value=0.0036), therefore pairwise treatment comparisons were performed (Table 4). Based on these results, there were significant differences (p-value=0.0030) between treated and untreated birds infected with the drug-susceptible isolate (Ow), and between treated birds infected with the susceptible (Ow) and resistant (Sn) isolates (p-value=0.0150). However, there were no significant differences (p-value=0.2600) between treated and untreated birds infected with the resistant isolate (Sn).

**Table 2.** Feed conversion ratio for each group by week. Feed conversion was calculated as kilograms of feed divided by weight gain.

| Week | Treatment |  |  |  |
| --- | --- | --- | --- | --- |
|  | Ow-Treated | Ow-Untreated | Sn-Treated | Sn-Untreated |
| 1 | 1.26 | 1.21 | 1.16 | 1.18 |
| 2 | 1.30 | 1.34 | 1.82 | 1.32 |
| 3 | 1.50 | 1.46 | 1.40 | 1.48 |
| 4 | 1.55 | 1.56 | 1.52 | 1.64 |
| 5 | 2.04 | 1.77 | 1.87 | 2.01 |
| 6 | 1.71 | 2.01 | 1.83 | 1.94 |
| 7 | 1.74 | 2.12 | 2.05 | 2.01 |
| 8 | 2.59 | 2.90 | 2.98 | 3.16 |
| 9 | 2.48 | 2.83 | 2.58 | 2.82 |

**Figure 1.**
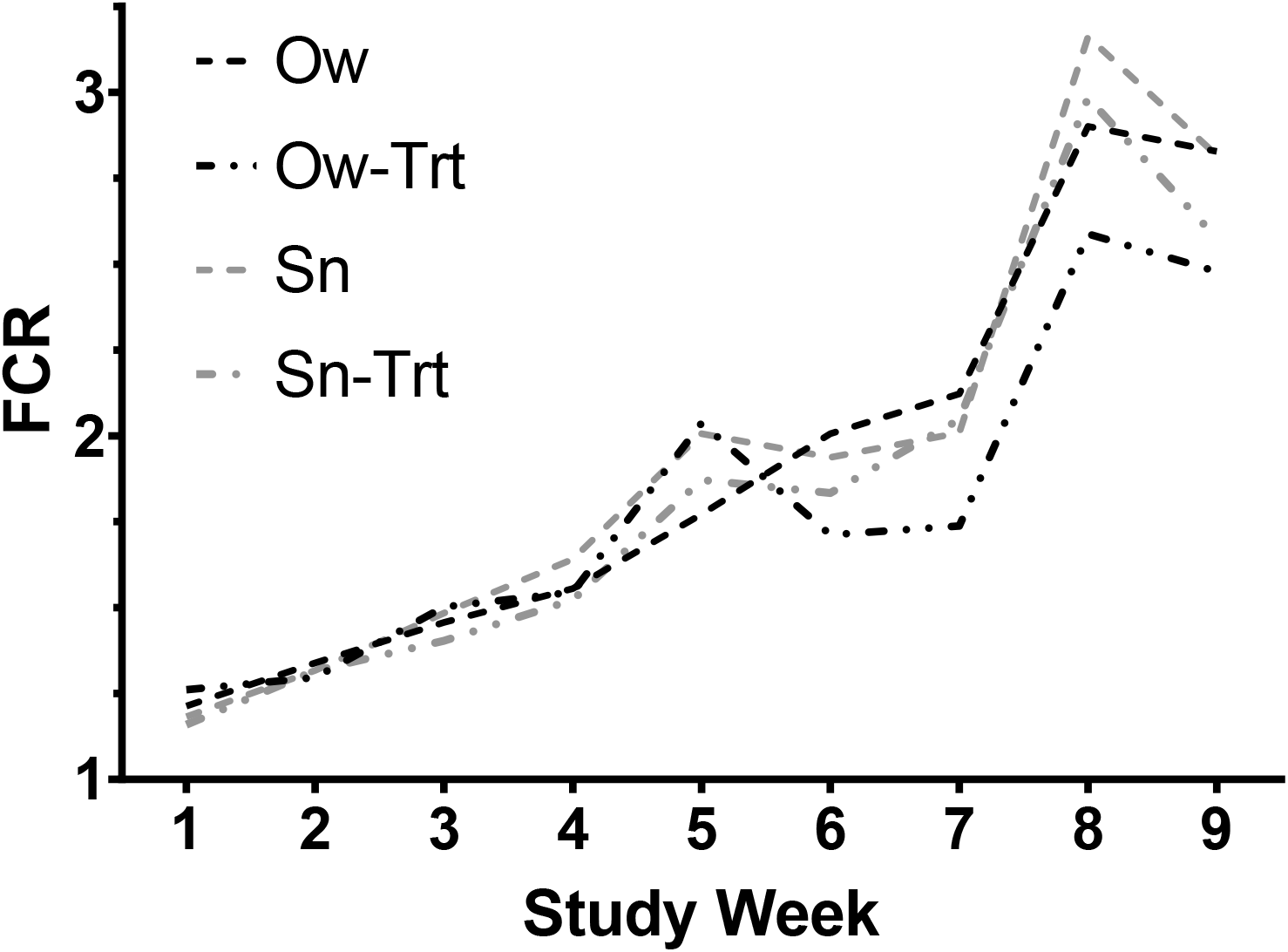
FCR for each treatment group over time.

**Table 3.** Least square means for FCR of each treatment group.

| Treatment | Estimate | Standard Error |
| --- | --- | --- |
| Ow-Treated | 0.7241 | 0.02995 |
| Ow-Untreated | 0.8645 | 0.02917 |
| Sn-Treated | 0.831 | 0.02762 |
| Sn-Untreated | 0.8755 | 0.02663 |

**Table 4.**
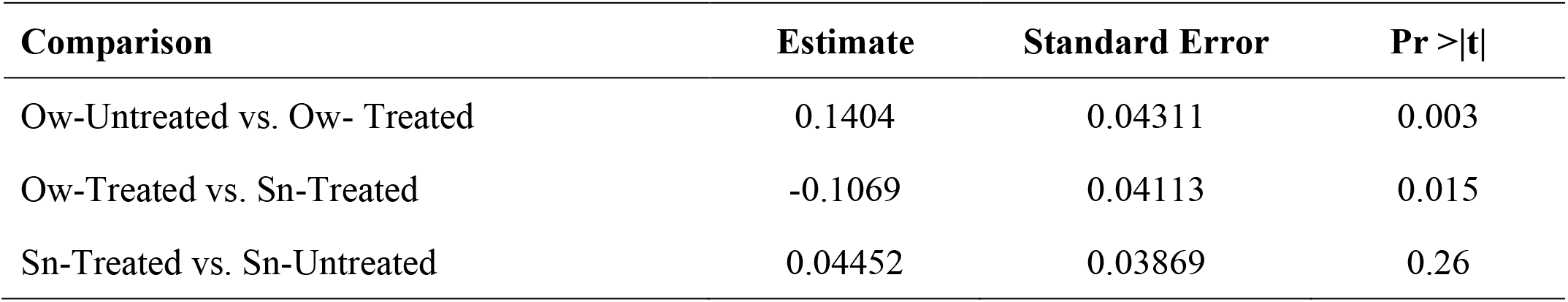
Pairwise comparisons for differences in least square means for FCR.

| Comparison | Estimate | Standard Error | Pr >\|t\| |
| --- | --- | --- | --- |
| Ow-Untreated vs. Ow- Treated | 0.1404 | 0.04311 | 0.003 |
| Ow-Treated vs. Sn-Treated | -0.1069 | 0.04113 | 0.015 |
| Sn-Treated vs. Sn-Untreated | 0.04452 | 0.03869 | 0.26 |

### Worm Counts at Week 7

The treated Ow group had no adults recovered, thus no analyses were performed for this group. No significant differences were seen between the treated and untreated Sn groups (p-value=0.8138). Additionally, there were no significant differences in adult worms between the untreated Ow group and the untreated Sn (p-value=0.4832) or between the untreated Ow group and treated Sn groups (p-value=0.2652). There were significant differences between the untreated and treated groups in the number of immature worms recovered for both the Ow (p-value = 0.0112) and Sn groups (p-value =0.0204). However, there were no significant differences between the treated Ow and treated Sn groups in the number of immature worms recovered (p-value = 0.1452). Mean worm counts for each treatment group at Week 7 are shown (Table 5).

**Table 5.** Mean worm counts by group at Week 7 and Week 9. For each treatment group, 8 birds were necropsied at week 7, and 16 birds were necropsied at week 9. Statistically significant groups are designated. No analysis was done on total worm burdens.

| **Group** | **Week 7** |  |  | **Week 9** |  |  |
| --- | --- | --- | --- | --- | --- | --- |
|  | **Immature** | **Adults** | **Total** | **Immature** | **Adults** | **Total** |
| Ow-Treated | 1.88<sup>c</sup> | 0.00<sup>b</sup> | 1.88 | 0.13<sup>b</sup> | 0.00<sup>b</sup> | 0.13 |
| Ow-Untreated | 4.50<sup>ab</sup> | 3.38<sup>a</sup> | 7.88 | 10.94<sup>a</sup> | 0.38<sup>a</sup> | 11.31 |
| Sn-Treated | 3.13<sup>bc</sup> | 2.25<sup>a</sup> | 5.38 | 8.38<sup>a</sup> | 1.06<sup>a</sup> | 9.44 |
| Sn-Untreated | 6.00<sup>a</sup> | 2.50<sup>a</sup> | 8.50 | 11.13<sup>a</sup> | 0.44<sup>a</sup> | 11.56 |

### Worm Counts at Week 9

Very few adult worms were recovered from any of the groups and most birds had no adult worms. Accordingly, no significant differences in adult worms were noted. There were, however, significant differences in the number of immature worms between the untreated and treated groups for both Ow birds (p-value <0.0001) and Sn birds (p-value <0.0001). Additionally, significant differences were observed in the number of recovered immature worms between the treated Ow group and treated Sn group of birds (p-value <0.0001). Mean worm counts for each treatment group at Week 9 are shown (Table 5).

## 4. DISCUSSION

To the best of our knowledge, here we report findings of the first study measuring the effects of drug-resistant *A. dissimilis* infection on turkeys. By infecting groups of birds with either a known fenbendazole-susceptible and known fenbendazole-resistant isolate, we were able to determine, using a mixed model for comparisons, the level of production loss caused by drug-resistant parasites which were not removed by treatment. This model allowed for comparisons that accounted for the random variability of worm burdens, feed consumption, etc. For these comparisons, results were analyzed using both week 4 and week 5 as a baseline and no differences in statistical results were seen using either week as baseline. Thus, we used week 5 as baseline for all comparisons, as this was the point from which measurements would begin to diverge as a consequence of failed treatments due to the presence of resistant worms.

Significant differences seen in FCR between the treated and untreated drug-susceptible Ow groups indicate that the *A. dissimilis* infections were impairing FCR, and successful removal of the drug-susceptible worms by treatment led to higher feed efficiency. In contrast, treatment of birds infected with the drug-resistant Sn isolate did not yield an improvement in FCR. Interestingly, no differences were seen in weight gain between groups, highlighting that this effect on FCR is solely on feed consumption. Feed conversion efficiency is significantly diminished, but birds appear to have gorged themselves on feed, making up for any possible weight loss and driving FCR higher. Beginning in week 6 through the end of the study, the treated Ow groups consumed an average of 230 grams less feed per week per bird as compared to the treated Sn groups.

If the levels of production loss seen in this study due to the drug-resistant worms were extended to the level of a house of 10,000 birds, this difference in feed usage would translate to an extra 2.3 metric tons of feed needed per week. Using our feed cost of approximately $275 US dollars/metric ton, this amounts to around $635 in extra feed costs/per week. Our grow-out only lasted for 9 weeks, thus projections for a full grow-out if 16 weeks need to be made cautiously. However, if this difference is projected onto a full 16 week grow-out, starting from week 5, total extra feed costs due to effects of *A. dissimilis* on FCR for a 10,000-bird house would be approximately $6,985.

The rather large differences recorded in FCR in this study are even more dramatic when viewed in light of the low worm burdens achieved in this study. In a previous study with *A. dissimilis* performed in commercial houses, mean worm burden from natural infections at day 56 post-infection was 13 adult worms per bird (Yazwinski, Rosenstein et al. 1993). In our recent study, mean worm burdens from a bolus infection administered by gavage averaged 18.3 adult worms per bird in untreated birds (Collins, Jordan et al. 2019). In contrast, at week 7 in our current study (49 days post-infection), our untreated groups had average adult worm burdens per bird of only 8.5 and 7.9 for Sn and Ow, respectively. This is only around 25% of what was seen in the Yazwinski study at a similar time point, and around 44% of the burden seen in our previous study. An estimated 200 total eggs per bird were given both in our previous, as well as the current study. In the present study, our infection protocol was designed to replicate the trickle infection birds would be expected to experience in a commercial house, however it failed to produce the worm burdens seen in these previous studies. Despite this, we were still able to determine the effects of treatment of worm burden in our treatment groups.

At week 7, in agreement with the significantly improved FCR, no adult parasites were recovered from necropsy of Ow-Treated birds, indicating the high efficacy of fenbendazole against this susceptible isolate by eliminating 100% of the adult burden. The few immature parasites recovered from this group are most likely due to reinfection in the intervening post treatment period. At this same time point, there were no significant differences in worm burdens between treated and untreated Sn groups, and both had significantly higher adult worm burdens than the treated Ow group, but not the Ow untreated group indicating the inability of treatment to control parasites of the resistant isolate. This lack of control is in agreement with the lack of improvement seen in FCR at this time point.

Although we were able to detect an impact on feed conversion, larger worm burdens more typical of natural infections are needed to determine the full scale of drug-resistant worms on FCR. It seems likely that higher worm burdens would have produced even greater negative impacts on FCR than what are reported here. In addition to burdens, rearing time likely also plays an important role in the effects on FCR. Longer grow out times with continual reinfection due to environmental contamination with infective eggs, may lead to heavier burdens and therefore increase the impacts. Due to limitations of our research space, which was designed for chickens, it was necessary to prematurely terminate the study after 9 weeks. This contrasts to the typical commercial grow out of 16-20 weeks. With a longer grow out period, it is possible that the effects on FCR would continue or worsen causing further costs associated with resistant parasites. Little is known about the population dynamics of *A. dissimilis*, and these dynamics, would likely play a large role in determining the effects of resistant parasites in a full grow-out. Additional studies will be needed to address this issue.

Overall, our data suggests that fenbendazole-resistant *A. dissimilis* have the potential to impart substantial economic losses in the production of commercial turkeys. Presently, the prevalence of resistance to fenbendazole is unknown, but may be much higher than is currently realized (Collins, Jordan et al. 2019). Taken together, the results of our two recent studies highlight the need for surveillance of resistance in helminths of poultry, for developing strategies to prevent the development of drug resistance, and for developing strategies to address the presence of drug resistant worms on a farm. Additional studies that better replicate the grow-out time and worm infection levels that are typical on commercial turkey farms are needed to gain a more accurate and full measure of the economic impacts of resistant *Ascaridia dissimilis* on turkey production.

## 5. CONCLUSION

This study highlights the fact that *A. dissimilis* can significantly impact the economy of turkey production even with low sub-clinical levels of infection. Thus, drug-resistant *A. dissimilis* have the potential to significantly impact the production economy of turkeys.

## ACKNOWLEDGEMETS

We would like to thank Dr. Justin Fowler for formulation the starter and grower ration. We would also like to thank Sue Howell, Dr. Kelsey Paras, Mikayla Bazemore, Kyle Harris, Patricia Weatherly, Dr. Cassan Pulaski, Will Van Brackle, Abbey Malatesta and Dr. Daniel Zuluaga for their assistance in multiple phases of this study.

This project was supported by a grant from the US Poultry and Egg Association (project #F081)

## ETHICAL STATEMENT

All birds were handled under protocols approved by the University of Georgia Institutional Animal Care and Use Committee (IACUC) under animal use policy A2019 01-005-Y2-A1.

## Notes

### Competing Interest Statement

The authors have declared no competing interest.

### Summary of Updates

Setting title and abstract genus/species names to Italics

