## Supplemental Tables 1 and 2 for "Impact of fenbendazole resistance in *Ascaridia dissimilis* on the economics of production in turkeys"

**Supplementary Table 1. Least square means for FCR, using Week 4 as Baseline**

| Treatment | Estimate | Standard Error |
|---|---|---|
| Ow-Untreated | 0.8172 | 0.02404 |
| Ow-Treated | 0.7173 | 0.02771 |
| Sn-Untreated | 0.8317 | 0.02318 |
| Sn-Treated | 0.7961 | 0.02382 |

**Supplementary Table 2. Pairwise treatment comparison for differences in least square means for FCR, using Week 4 as baseline, between Ow-Untreated and Ow-Treated, Ow-Treated and Sn-Treated, and Sn-Treated and Sn-Untreated.**

| Comparison | Estimate | Standard Error | Pr >\|t\| |
|---|---|---|---|
| Ow-Untreated vs. Ow- Treated | 0.09983 | 0.03666 | 0.0112 |
| Ow-Treated vs. Sn-Treated | -0.07875 | 0.03639 | 0.0395 |
| Sn-Treated vs. Sn-Untreated | 0.03565 | 0.0338 | 0.3009 |
